# Development of Pavlovian-to-instrumental transfer (PIT) task to examine the effects of reward-predicting cues on behavioral activation in young adults

**DOI:** 10.1101/601195

**Authors:** Raquel Quimas Molina da Costa, Emi Furukawa, Sebastian Hoefle, Jorge Moll, Gail Tripp, Paulo Mattos

## Abstract

There is a growing recognition that much of human behavior is governed by the presence of classically conditioned cues. The Pavlovian-to-Instrumental Transfer (PIT) paradigm offers a way to measure the effects of classically conditioned stimuli on behavior. In the current study, a novel PIT paradigm was developed for use in conjunction with an fMRI classical conditioning task. This could allow for a measurement of BOLD responses to anticipated rewards, unconfounded by operant responses, while providing a behavioral measure of conditioning effects. Twenty-four healthy young adults completed 1) instrumental training, 2) Pavlovian conditioning, and 3) Transfer test. During instrumental training, participants learned to apply force on a handgrip to win money from slot machines pictured on a computer screen. During Pavlovian conditioning, slot machines appeared with one of two abstract symbols (cues), one symbol was predictive of monetary reward. During the Transfer test, participants again applied force on a handgrip to win more money. This time, slot machines were presented with the Pavlovian cues, but with the outcomes hidden. The results indicated increased effort on the instrumental task, i.e., higher response frequency and greater force, in the presence of a reward-predicting cue. Our findings add to a growing number of studies demonstrating PIT effects in humans. This paradigm was proved to be useful in measuring the effects of a conditioned stimulus on behavioral activation.

## Introduction

Pavlovian, or classical, conditioning refers to an associative process through which neutral stimuli acquire motivational significance after repeated pairing with a rewarding or aversive experience (Corbit and Balleine, 2003). When a previously neutral stimulus is repeatedly paired with reward, the stimulus comes to elicit physiological responses and induce appetitive behaviors that are normally associated with reward (Cartoni, Balleine and Baldassarre, 2016). Such appetitive classical conditioning is thought to underlie much of everyday human behavior (Bray *et al*., 2008; Cartoni, Balleine and Baldassarre, 2016), helping maintain both adaptive and maladaptive behavior, for example addiction (Hogarth, 2012; Garbusow *et al*., 2016). To date, only a small number of experimental studies have shown clear behavioral effects of appetitive conditioning in humans (Vogel *et al*., 2018). This is in contrast to substantial literature documenting the effectiveness of aversive conditioning (Mineka and Oehlberg, 2008; Maren, Phan and Liberzon, 2013; Rigoli, Pezzulo and Dolan, 2016; Mkrtchian, Roiser and Robinson, 2017).

In animal experiments, effects of appetitive classical conditioning are studied using primary rewards, such as food, which elicits innate, reflexive responses (Corbit and Balleine, 2011). Providing primary rewards repeatedly in experiments with human subjects can be challenging and deprivation prior to an experiment (e.g., hunger or thirst) may be required to ensure reward saliency (Freeman *et al*., 2015; Manglani *et al*., 2017; Schad *et al*., 2018). When well-established secondary rewards (such as images of food, money or points) are used in the place of primary rewards, the consumption of rewards is typically delayed (e.g., accumulating points during but receiving actual rewards after an experiment). Such procedural constraints can interfere with observation of simple behavioral activation by classically conditioned cues, e.g., approach behavior to the food hopper in animal experiments.

The Pavlovian-to-Instrumental Transfer (PIT) paradigm offers a way to measure the effects of conditioned stimuli on behavior in humans. In a PIT task, the motivational influence of reward-predicting cue is measured through their effects on independently learned instrumental behavior (Cartoni, Balleine and Baldassarre, 2016). A PIT task has three components. During Pavlovian conditioning, subjects are exposed to previously neutral cues (e.g., sounds), at least one of which is repeatedly followed by rewards (e.g., food). In a separate instrumental conditioning phase, subjects are trained to engage in behavior (e.g., lever press) to obtain rewards. During the transfer phase, Pavlovian cues are presented while subjects have the opportunity to engage in the previously learned instrumental behavior. If the cues have acquired motivational value, their presence should invigorate the behavioral response. The transfer phase is carried out under extinction, so that instrumental behavior is not modified by ongoing rewards (Levey and Martin, 1991; Corbit and Balleine, 2011).

A number of human PIT studies has been reported in recent years, reflecting growing interest in understanding the effects of Pavlovian cues on behavior. A little more than half of the studies examined appetitive conditioning. Many of these studies have focused on examining addictive behavior, e.g., to food (Pool *et al*., 2015; Quail, Laurent and Balleine, 2017; Seabrooke *et al*., 2017; van Steenbergen *et al*., 2017), to nicotine (Hogarth, 2012; Hogarth, Maynard and Munafò, 2015; Manglani *et al*., 2017) and to alcohol (Martinovic *et al*., 2014; Garbusow *et al*., 2016; Hardy *et al*., 2017; Sommer *et al*., 2017a). Other studies have used PIT paradigms to evaluate how stress and depressed mood affect motivation (Huys *et al*., 2016; Quail, Laurent and Balleine, 2017), or to study the neural correlates of the transfer effects in non-disordered populations (Paredes-Olay *et al*., 2002; Talmi *et al*., 2008a; Allman *et al*., 2010; Geurts *et al*., 2013; Hebart and Gläscher, 2015; Sebold *et al*., 2016).

In many human appetitive PIT studies, the transfer is examined through the effects of conditioned stimuli on goal-oriented behavior, most often response frequency or go/no-go response accuracy (Garbusow *et al*., 2014a; Sebold *et al*., 2016). Specific transfer effects are most frequently documented, i.e., the presence of a conditioned stimulus motivates only the behavior previously associated with the same reward, e.g., increased responding on a specific choice, (Manglani *et al*., 2017; van Steenbergen *et al*., 2017). General invigoration of behavior appear more difficult to demonstrate in humans. We identified only five appetitive PIT studies that report general transfer effects, i.e., increased responses in the presence of a conditioned stimuli, specific rewards associated with which were not available during instrumental training (Prevost *et al*., 2012a; Lovibond and Colagiuri, 2013; Hebart and Gläscher, 2015; Huys *et al*., 2016; Quail, Laurent and Balleine, 2017).

Efforts to understand the effects of appetitively conditioned stimuli in humans extend to fMRI studies. A small number of studies used simple classical conditioning paradigms to study the neural correlates of reward-predicting cues (e.g., Prevost *et al*., 2012b; Furukawa *et al*., 2014; Klucken *et al*., 2016). In these studies, as no behavioral response was required, conditioning effects on behavior were not measured. In other studies, participants were instructed to make a behavioral response following a reward-predicting cue, with fast accurate responding a measure of subjects’ motivation (e.g., Knutson *et al*., 2000; O’Doherty, Hampton and Kim, 2007). In these paradigms, it is difficult to determine if observed BOLD responses reflect anticipation of acting to obtain reward or anticipation of the reward itself.

We developed a novel PIT paradigm, for use in conjunction with fMRI classical conditioning tasks. The paradigm facilitates separation of BOLD responses to anticipated actions and anticipated rewards, while providing a behavioral measure of classical conditioning effects. Instrumental behavior was established prior to fMRI scanning; participants learned to grip a hand dynamometer to obtain monetary rewards. Classical conditioning took place in the MRI scanner. A previously neutral cue was paired with monetary reward outcome, another cue with no-reward outcome. Transfer effects, handgrip responses in the presence of a now reward-predicting cue, were evaluated following the scanning session. In this paper, we describe the PIT paradigm in detail together with the behavioral effects of the reward-predicting cue.

## Methods

Ethical approval for the study was obtained from the ethics committee of the D’Or Institute for Research and Education (IDOR), Brazil. All participants were volunteers and provided written consent.

### Participants

The study included data from 24 typically developing young adults (Mean age = 27 ± 3.55, Mean Estimated IQ = 104 ± 5.79, Mean education level = 17 ± 1,77, 54.16 % female), recruited at local universities and through IDOR researchers’ personal contacts. All participants belonged to middle and upper socioeconomic classes (http://www.abep.org/). Participants completed a demographic and background questionnaire, an abbreviated IQ test (Wechsler Abbreviated Scale of Intelligence Vocabulary and Block Design subtests (Weschler, 1999)) and structured interviews with a psychiatrist (Kiddie-Schedule for Affective Disorder and Schizophrenia-PL (Brasil, 2003) and Structured Clinical Interview (Del-Ben *et al*., 2001)). The inclusion criteria for the study were: no major depressive or bipolar disorder, attention-deficit hyperactivity disorder, neurological disorder, current drug use or psychotic symptoms. Individuals with conditions associated with altered motivational processes, i.e., depressed or elevated mood, hyperactivity and impulsivity symptoms, addictive behavior, were excluded (Tripp and Wickens, 2008; Whitton, Treadway and Pizzagalli, 2015; Conzelmann *et al*., 2016).

### Experimental task

The PIT paradigm included Instrumental training, Pavlovian conditioning and a Pavlovian-to-Instrumental Transfer test. The task was programmed using Presentation^®^ version 16.5, by Neurobehavioral Systems Inc. Participants completed the Pavlovian conditioning inside a 3T Achieva scanner (Philips Medical Systems, The Netherlands), with task stimuli shown using an LCD display and mirror adapted to the head coil. The instrumental training and transfer test were conducted outside the MRI scanner with the stimuli presented on a computer screen. Instrumental responses were measured using a hand dynamometer (www.biopac.com) calibrated with a BIOPAC MP160 System with AcqKnowledge software (Biopac^®^ Systems Inc). Participants were told they would receive their winnings (monetary rewards) accumulated throughout the experiment in the form of a gift card.

Before instrumental training, each participant’s maximum grip force was measured. Participants were asked to grip the dynamometer as hard as possible three times, from these grips their mean maximum force was calculated. Participants were instructed to grip the hand dynamometer to spin one of two “slot machines” pictured side by side on a computer screen (Figure 1). They practiced spinning a slot for 10 seconds. Each machine was programed to start spinning when the grip force reached 40% or 60% of the participant’s maximum force, requiring them to exert effort. The force required alternated semi-randomly to prevent participants modulating their grip to a set level.

**Figure 1.**
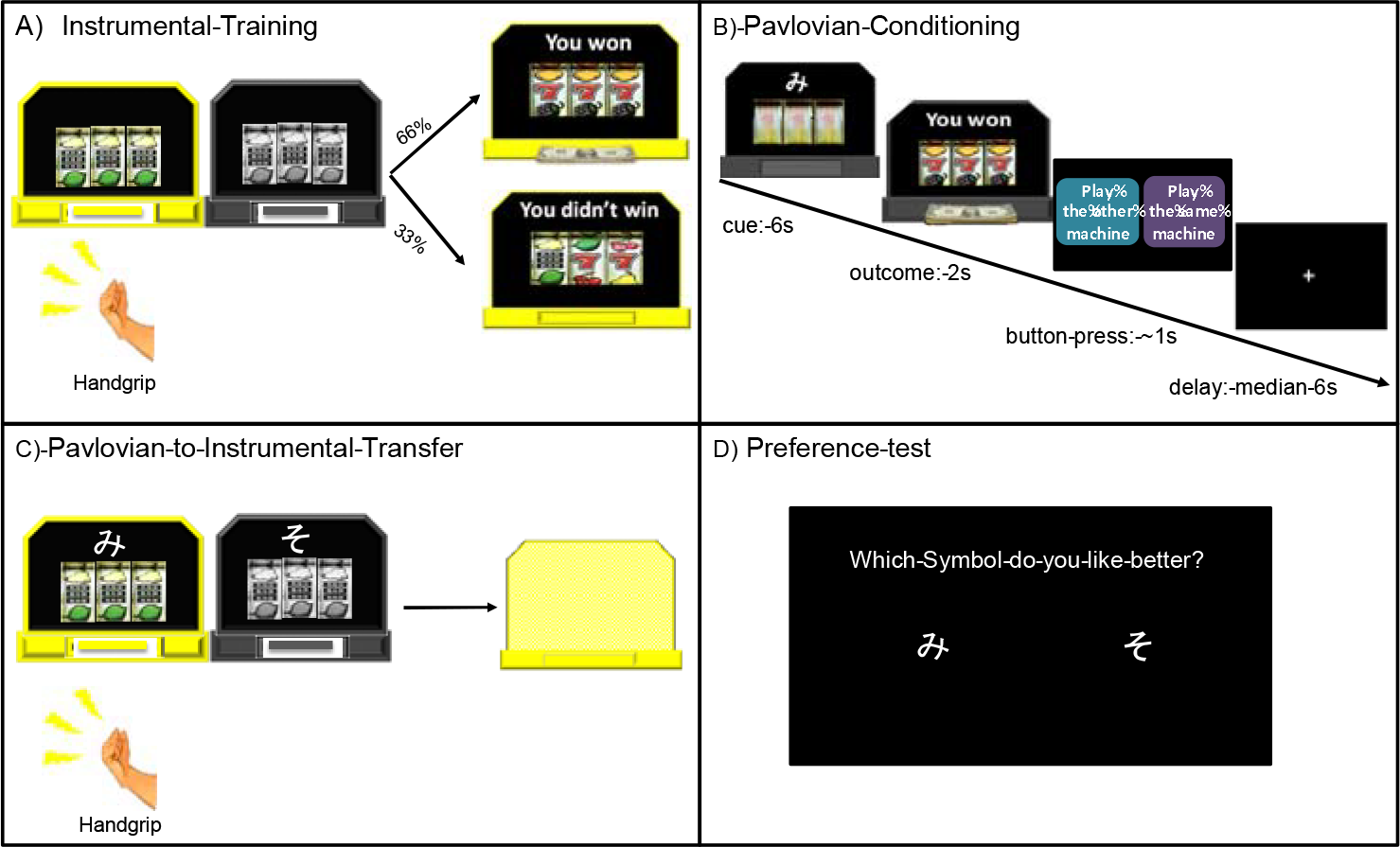
Pavlovian-to-Instrumental Transfer paradigm.

At the start of instrumental training the two slot machines were both gray. One machine at a time lit up and became available for a participant to play for 10 seconds before switching off. The two machines were identical during instrumental training, during the Transfer test each machine was shown with a different symbol (cue). The participants were told they could grip as many times as they wanted while a machine was lit and the more times they played the more chances they had to win. There was a 66% chance of winning on either machine. There were 12 ten second blocks with five second rest periods between each block. If participants failed to apply the necessary force to spin the slot for three consecutive seconds, the block ended. Machine availability alternated in a semi-random order over the 12 blocks, with all participants playing each machine for six blocks. The instrumental training lasted for approximately three minutes.

During Pavlovian conditioning, participants were told to watch the computer play the slot machines, and that the earnings would be added to their gift card. One of two abstract symbols (Cue A and Cue B; two Japanese hiragana letters) was displayed on top of each slot machine. Machine A (with Cue A) yielded reward 66% of the time (CS+). Machine B (with Cue B) was never associated with winning (CS−). The slot machine spun for six seconds, and an outcome was displayed for two seconds. After each trial, participants were asked to ‘suggest’ to the computer whether to play the same machine (with the same cue) or the alternative machine by pressing one of two buttons placed in their right hand, within a one second response window. This “choice” was designed to maintain participants’ attention during conditioning and did not influence machine presentation. Actual presentation order was semi-random with the constraint of no more than four consecutive trials with the same cue, and was the same for all participants. Pavlovian conditioning lasted for approximately 27 minutes and included 44 trials of Cue A (CS+) with reward outcome, 22 trials of Cue A with non-reward outcome, and 44 trials of Cue B (CS−). At the end of this phase, participants were asked which of the two symbols (cues) they preferred.

The transfer test was identical to instrumental training with the following exceptions: a) the reward and non-reward cues from the Pavlovian conditioning phase were displayed with their associated slot machines and b) the outcomes were “hidden” from the participants to allow examination of the behavioral effects of the cues in the absence of ongoing contingencies. Participants were told the computer would record their earnings, which would be added to their gift card. There were 12 ten second blocks for all participants: six blocks with Machine A (CS+) and six blocks with Machine B (CS−), becoming available in a semi-random order. At the end of the transfer test, the total money earned during the experiment was displayed for the participants on the screen (total amount = R$ 0.33 x the number of wins). All participants received gift cards with the total amount earned, which ranged from R$30 to R$50.

### Data analysis

To obtain a baseline measure of the participants’ instrumental learning, the frequency of grips above the maximum force threshold and the force applied (including the grips below the threshold) were recorded for each of the two slot machines. During the transfer test, the frequency of grips and the force levels were again recorded. Paired t-tests were used to compare if these performance measures differed between the two machines (CS+ vs. CS−). In addition, block effects for the instrumental training and transfer test were analyzed using repeated-measures GLM. Participants’ responses during the Pavlovian phase were also examined to assess their preference toward CS+ (log10[percent stay after CS+/(1 - percent stay after CS+)]) versus CS-(log10[percent stay after CS-/(1 - percent stay after CS−)]). Percentages of missed response window after three trial types, i.e., CS+ followed by reward, CS+ followed by non-reward, and CS-followed by non-reward, were also compared.

## Results

### Instrumental training

All participants learned to apply force to the handgrip to spin the slot machine during instrumental training. We did not expect that participants’ behavior would be different across the two slot machines. This was confirmed by paired t-tests that showed no difference in the mean force level applied and frequency of grips between the two machines. The frequency of grips per block declined over the 12 blocks, F(3.56,81.89)= 3.05, p < .05, Greenhouse-Geisser correction, likely reflecting fatigue. The mean force remained stable suggesting continued engagement in the task throughout the phase.

### Pavlovian conditioning phase

A paired sample t-test showed a greater proportion of suggested stays following the CS+ (88%) versus CS-(14%) trials, t(23) = 8.25, p < .001. Moreover, the participants were significantly more likely to suggest staying on the same machine following CS+ non-reward trials (80%) than CS-non-reward trials CS-(14%), t(23) = 5.72, p < .001. This indicates the responses were made based on the cues.

All participants responded, i.e., suggested which machine to play next, during the one second window more than 75% of the time, indicating adequate task attention. Participants were less likely to miss the response window when the CS+ was followed by reward outcome (7%) than non-reward outcome (54%), t(23) = −15.32, p < .001. Similarity, they were also less likely to miss responding after CS-trials (all non-reward outcome) (8%), than after CS+ non-reward trials (54%), t(23) = −14.54, p <.001.

All participants indicated that they preferred Cue A (CS+) over Cue B (CS−) when questioned after completing Pavlovian conditioning.

### Pavlovian-to-instrumental transfer phase

Paired sample t-tests indicated that participants applied greater force, t(23) = 5.06, p < .001, and responded more frequently, t(23) = 7.40, p < .001, on the Machine A (displayed with CS+) than the Machine B (displayed with CS−) (Table 1). The GLM examining performance over the 6 blocks for each cue type also showed differential behavior effects of the conditioned cues. In the presence of the CS+, participants maintained the frequency of grips (per block) over time. Response frequency declined over the 6 blocks in the presence of the CS−, F(3.06,70.26) = 5.87, p < .01, Greenhouse-Geisser correction. The mean force was maintained over six blocks for both machines.

**Table 1.** Response frequency and force during the instrumental and transfer phase.

|  | Response Frequency |  |  |  |  | Grip Force (kg) |  |  |  |  |
| --- | --- | --- | --- | --- | --- | --- | --- | --- | --- | --- |
| Instrumental | Slot machine 1 |  | Slot machine 2 |  |  | Slot machine 1 |  | Slot machine 2 |  |  |
|  | Mean | sd | Mean | sd | Paired t test<br>(p value) | Mean | sd | Mean | sd | Paired t test<br>(p value) |
|  | 6.68 | 1.22 | 6.63 | 1.23 | 0.43 (.674) | 3.19 | 1.35 | 3.22 | 1.33 | -0.81 (.427) |
| Transfer | Response Frequency |  |  |  |  | Grip Force (kg) |  |  |  |  |
|  | Slot machine A<br>(CS+) |  | Slot machine B<br>(CS-) |  |  | Slot machine A<br>(CS+) |  | Slot machine B<br>(CS-) |  |  |
|  | Mean | sd | Mean | sd | Paired t test<br>(p value) | Mean | sd | Mean | sd | Paired t test<br>(p value) |
|  | 6.80 | 1.11 | 2.13 | 2.79 | 7.40 (.000*) | 2.89 | 1.16 | 1.52 | 1.24 | 5.06 (.000*) |

## Discussion

The current study evaluated the effects of reward-predicting cues on behavior, using a PIT task in which classical conditioning trials were optimized for fMRI. The PIT task included three phases; instrumental training, classical (Pavlovian) conditioning, and a transfer test. The behavioral effects of appetitive conditioning were demonstrated with this new PIT task. The presence of a conditioned stimulus motivated individuals to engage in effortful behavior.

During instrumental training, participants learned to apply force to a hand dynamometer to spin the slot machines to earn monetary rewards. Under Pavlovian conditioning, participants demonstrated a behavioral preference for the slot machine associated with the reward-predicting cue. Following conditioning, all participants selected the symbol coupled with reward when asked for their preference. During the transfer test, in the presence of the reward-predicting cue, participants applied more force to and gripped the hand dynamometer more frequently. The number of grips was stable across the response blocks in which the reward-predicting cue was present. Grip frequency declined over blocks in the presence of the non-reward cue. Response frequency was observed to decline over time during instrumental training when no cues were presented.

Our results are consistent with a growing number of studies demonstrating PIT effects in humans (Seabrooke *et al*., 2017; Sommer *et al*., 2017b; Verhoeven, Watson and de Wit, 2018; Vogel *et al*., 2018) and provide additional evidence for behavioral invigoration in presence of a classically conditioned stimulus. The current PIT paradigm was developed to allow Pavlovian conditioning to be completed in an MRI scanner. The Pavlovian conditioning phase was designed to ensure: 1) sufficient presentations of each cue-outcome pair and 2) that BOLD effects were not influenced by operant behavioral responses or complex reward probability or magnitude estimation. During the instrumental and transfer phases, grip strength, in addition to grip frequency, was recorded to quantify physiological effort. Motivation, as expressed in behavior, has been characterized as engaging in effortful action to obtain desirable outcomes (Bortolini *et al*., no date; Chong, Bonnelle and Husain, 2016).

Study limitations should be considered in interpreting the findings. All participants explicitly indicated a preference for the CS+ over the CS-; whether transfer effects would have been observed without such cognitive awareness is not known. Some studies have demonstrated awareness of the reinforcement contingencies affecting transfer effects (Lovibond and Shanks, 2002; Talmi *et al*., 2008b; Jeffs and Duka, 2017). We used a single CS+ to examine the behavioral effects of a reward-predicting cue. As a consequence, general vs. specific transfer effects could not be differentiated in this study. Transfer effects were evaluated as the difference in behavior activation in the presence of CS+ vs. CS-, than as a change from an active baseline with a neutral cue. Thus, the observed effects may be due to CS+ energizing and/or CS-inhibiting behavior (Quail 2017). Even in a simple PIT paradigm, in human studies, we are likely measuring some degree of explicit behavioral activation, informed by the conditioned cue, i.e., a decision to grip forcefully, or not. However, the maintenance of effortful behavior in the presence of the CS+ argues for its energizing effects.

The study findings indicate this new paradigm is useful in measuring the effects of a previously-neutral, reward-predicting cue, on behavioral activation. The simplicity of the paradigm allows for its use in fMRI studies, and with a variety of participants, including clinical populations (Garbusow *et al*., 2014b; Hogarth, Maynard and Munafò, 2015; Huys *et al*., 2016). Altered sensitivity to reward-predicting cues has been hypothesized to contribute to a range of psychiatric such as addiction, ADHD and depression. Better understanding of the effects of Pavlovian cues on human behavior may help refine treatment strategies for people affected by these disorders.

## Acknowledgements

We would like to thank all individuals who took part in the study and IDOR clinical research and technical teams who assisted with data collection and processing.

